# The Bundles of Intercrossing Fibers of the Extensor Mechanism of the Fingers Greatly Influence the Transmission of Muscle Forces

**DOI:** 10.1101/2023.11.06.565148

**Authors:** Anton A Dogadov, Francisco J Valero-Cuevas, Christine Serviere, Franck Quaine

## Abstract

The extensor mechanism is a tendinous structure that plays an important role in finger function. It transmits forces from several intrinsic and extrinsic muscles to multiple bony attachments along the finger via sheets of collagen fibers. The most important attachments are located at the base of the second and third phalanges (proximal and distal attachments, respectively). How the forces from the muscles contribute to the forces at the attachment points, however, is not fully known. In addition to the well-accepted medial and lateral bands, there exist two layers of intercrossing fiber bundles (superficial interosseous medial fiber layer and deeper extensor lateral fiber layer), connecting them. In contrast to its common idealization as a minimal network of distinct strings, we built a numerical model consisting of fiber bundles to evaluate the role of multiple intercrossing fibers in the production of static finger forces. We compared this more detailed model of the extensor mechanism to the idealized minimal network that only includes the medial and lateral bands. We find that including bundles of intercrossing fibers significantly affects force transmission, which itself depends on finger posture. In a mid-flexion posture (metacarpal joint MCP = 45°; proximal interphalangeal joint PIP = 45°; distal interphalangeal joint DIP = 10°) the force transmitted by the lateral fibers is 40% lower than in a more pronounced flexed posture (MCP = 90°; PIP = 90°; DIP = 80°). We conclude that the intercrossing fiber bundles — traditionally left out in prior models since Zancolli’s simplification — play an important role in force transmission and variation of the latter with posture.

## INTRODUCTION

The extensor mechanism of the fingers of human and non-human primates is a network of tendinous structures that drapes over the dorsum of the finger bones (Van Zwieten, 1980). It transmits forces from several extrinsic and intrinsic hand muscles to the phalanges to produce torques at the finger joints (Landsmeer, 1949). This structure plays an important role in finger function, and its disruption degrades manipulation ability. Therefore, it is usually included in detailed biomechanical models of the fingers (Hu et al., 2014; Jadelis et al., 2023; Sachdeva et al., 2015; Valero-Cuevas et al., 2007; Vaz et al., 2015). Even though the extensor mechanism is, in reality, a sheet of intersecting fibers, it has often been idealized as a sparse network of strings (Chao, 1989; Garcia-Elias et al., 1991; Schultz et al., 1981; Valero-Cuevas et al., 2007; Zancolli, 1979). However, the extensor mechanism is a sophisticated continuous fibrous composite structure that can be simplified as having

1. A medial band, which originates from the extrinsic *extensor digitorum communis* muscle and has its principal bone insertion at the proximal part of the middle phalanx as the proximal band (Harris and Rutledge, 1972), *i*.*e*. proximal extensor mechanism attachment;
2. Two lateral (or intrinsic) bands, radial and ulnar, which originate from the intrinsic muscles. The radial and ulnar bands combine and insert to the proximal part of the distal phalanx as the terminal (Harris and Rutledge, 1972), *i*.*e*. distal extensor mechanism attachment;
3. The intercrossing fiber bundles and the extensor hood, connecting the lateral bands with the medial one (Schultz et al., 1981) The intercrossing fiber bundles are represented by two layers of fibers: interosseous medial fibers and the extensor lateral fibers.

The intercrossing fibers and the extensor hood are of particular interest because they biomechanically couple the forces in the medial and terminal bands and the rotations of both interphalangeal joints (Leijnse and Spoor, 2012). Moreover, the intercrossing fibers may become more tight or slack as a function of the posture (Leijnse and Spoor, 2012), making the force transmission among the extensor mechanism bands posture dependent (Lee et al., 2008; Sarrafian et al., 1970). This biomechanical coupling has been interpreted as also enabling a nonlinear transmission of tendon forces (i.e., a “switch” behavior) that improves controllability under the anatomical constraints that the fingers do not have any muscles in them (Valero-Cuevas et al., 2007). This means that changing the ratio between the input forces from the intrinsic and extrinsic muscles itself changes the distribution of forces across the proximal and terminal bands. However, we lack detailed studies identifying the posture-dependent interactions by which the multiple fiber bundles of the extensor mechanism enables finger function.

The purpose of this study is to fill this gap in understanding by using a more detailed model of the fiber bundles of the extensor mechanism to understand the role of the extensor hood and the intercrossing fibers on muscle force transmission to produce static fingertip force. In the current study, we focus, without loss of generality, on the extensor mechanism of the middle finger. Applied to the middle finger, the intrinsic muscles, mentioned above, are the second and the third dorsal interosseous muscles, and the second lumbrical muscle. In particular, we built and compared two three-dimensional models of the extensor mechanism: a more detailed model that includes the intercrossing fibers and an extensor hood, and a trivial model, without any structures connecting the central band with the lateral bands. We call it the “trivial” model because it reflects the theoretical baseline architecture of muscles where tendons originate in a muscle and insert into bone. While we do not endorse such a trivial structure, this trivial model is not a straw man. Rather, it is the baseline musculotendon anatomy, which evolutionary pressures—presumably of biomechanical nature—drove to specialize into an extensor mechanism. As such, it does help highlight and quantify the biomechanical benefits of a sophisticated extensor mechanism where tendons that originate in muscle combine with other tendons to then insert into bone.

Our results demonstrate changes in force transmission with changes in posture, introduced by the extensor hood and the intercrossing fiber bundles. The functional differences compared to the trivial model speaks to the evolutionary pressures that may have driven the evolution of the topology of the extensor mechanism in the first place, given the anatomical constraints that the fingers do not have any muscles in them and must be actuated by muscles in the palm and forearm. Our model simulating muscle force transmission via bundles of intercrossing fibers now allows us to better understand neuromuscular strategies for finger control, and explain the functional deficits associated with clinically common ruptures or adhesions of the elements of the extensor mechanism. It also enables the design of prostheses and robotics hands using such interconnected tendon architectures.

## METHODS

We coded a custom numerical environment that allows representing the extensor mechanism as contacting bundles of interconnecting fibers. Each string or consists of a sequence of points, pairwise connected by elastic elements with a linear stress-strain model. This computational environment is written using Matlab 2015 and C++, and is based on the extensor mechanism simulator, described in detail elsewhere (Dogadov et al., 2017). This environment allows simulating tendinous structures with arbitrary topologies and finger postures for static analysis. For a given vector of input muscle forces, it calculates the resulting net joint torques and fingertip wrench (endpoint forces and torques).

The first model (Fig. 1a) was a full extensor mechanism model that includes multiple bundles of intercrossing fibers to approximate the known anatomical bands and sheets of collagenous tissue. The model contains medial band (5), connecting the extrinsic *extensor digitorum* muscle with the proximal band (6). The latter forms a proximal attachment of the extensor mechanism to the skeleton. The model also contains the lateral (or interosseous) bands (4), connecting the intrinsic muscles with terminal band (9). The latter forms a distal attachment of the extensor mechanism to the skeleton. The attachment points of the tendons and ligaments to bones are shown by circles. Finally, the model contains the structures connecting the lateral band with the medial one. These structures are the extensor hood (1) and the bundles of intercrossing fibers: the interosseus medial fibers (2, shown in red) and the extensor lateral fibers (3, shown in blue). These bundles are shown enlarged in Fig. 2.

**Fig. 1.**
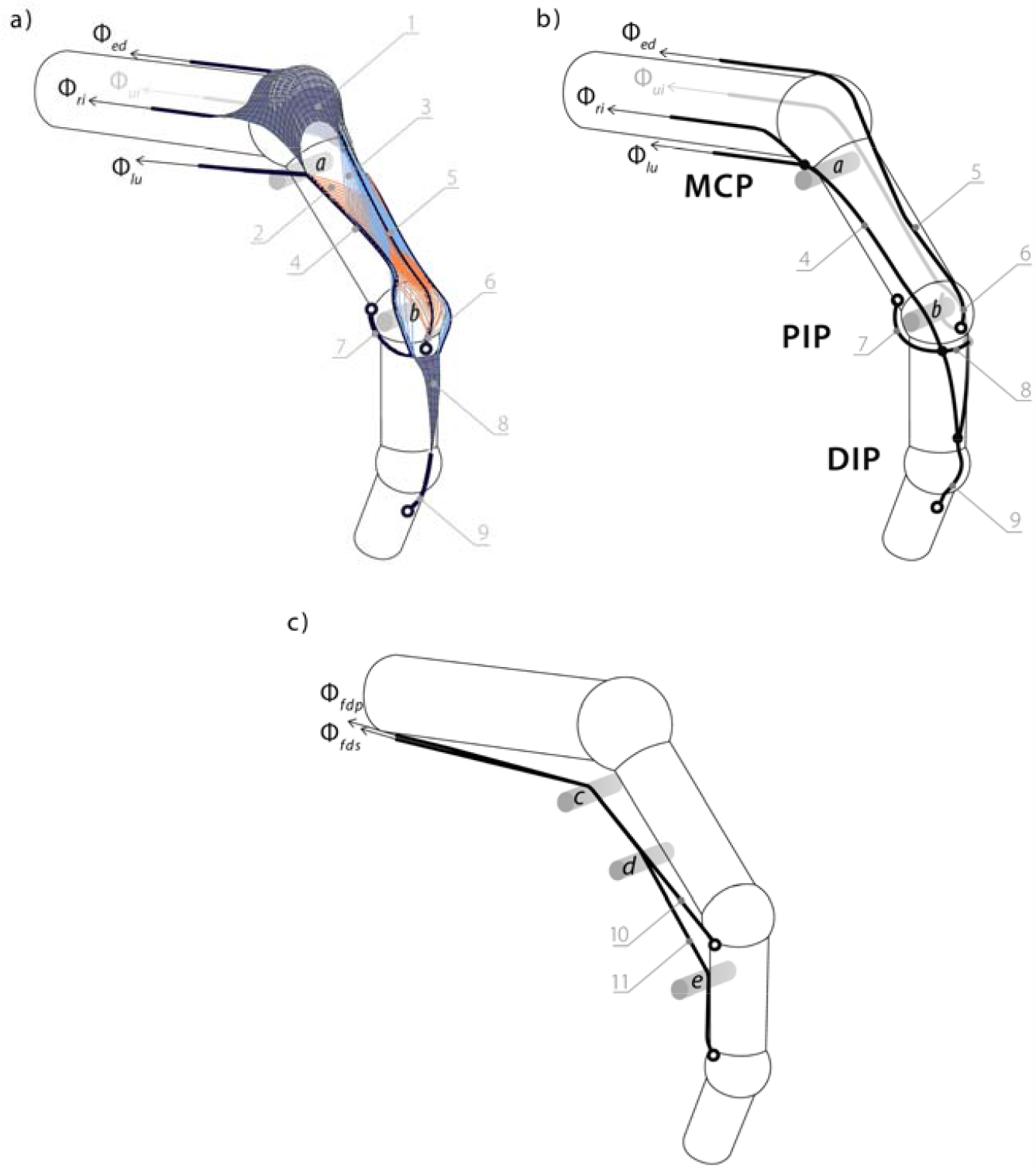
The view of the extensor mechanism modelled in a developed environment. a: the full model, which contains the principal tendon and ligaments of the extensor mechanism: 1 – the extensor hood, 2 – interosseous medial fibers (red), 3 – the extensor lateral fibers (blue), 4 – lateral band, 5 –medial band, 6 – proximal band, 7 – transverse retinacular ligament, 8 – triangular ligament, 9 – terminal band. b: the trivial model. The trivial model does not contain the structures connecting the lateral bands (4) with the extensor medial band (5). c: flexor tendons: 10 – flexor flexor digitorum superficialis tendon, 11 – flexor digitorum profundus tendon (same for both models)

**Fig. 2.**
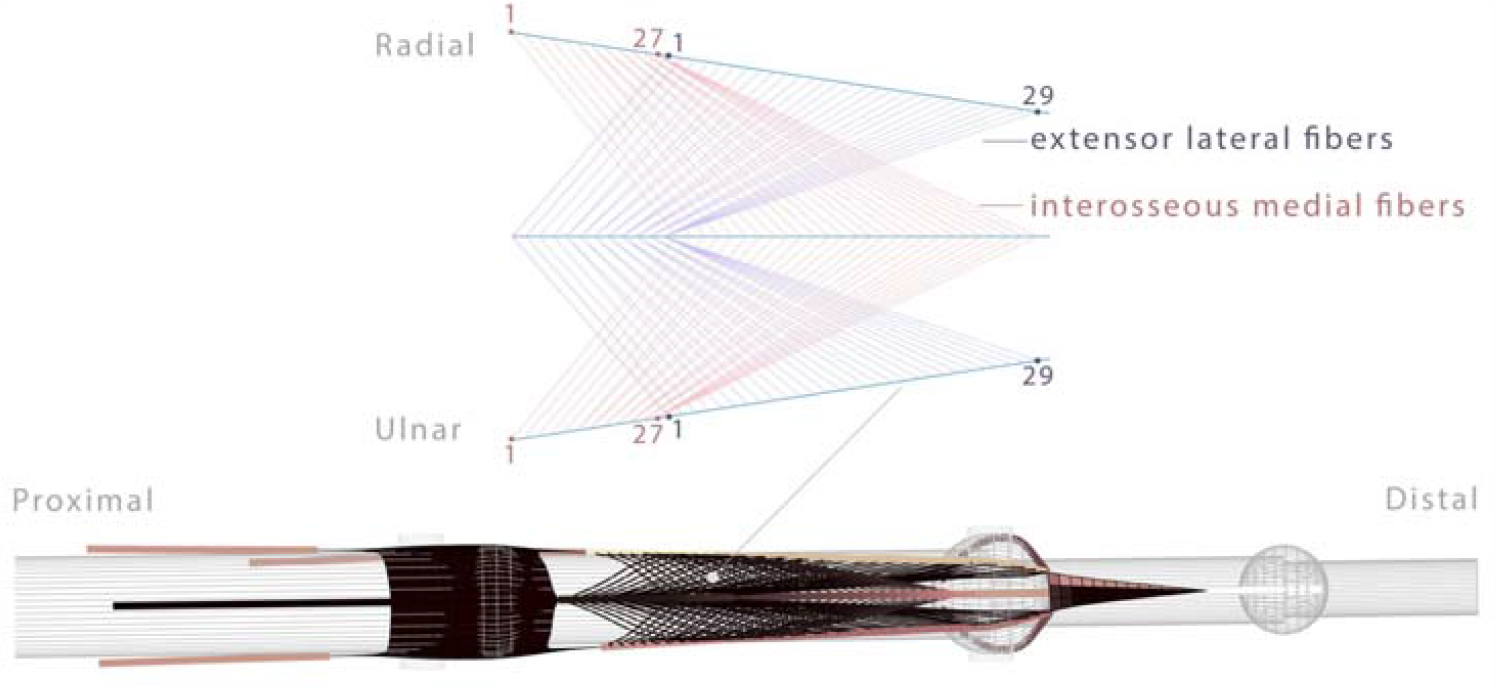
The schematic view of the intercrossing fibers. Red: interosseous medial fibers; blue: extensor lateral fibers.

The second model was the baseline (trivial) one (Fig. 1b), with no structures, connecting the lateral tendons with the medial one, *i*.*e*. it does not contain the extensor hood and intercrossing fiber bundles. The transverse retinacular ligament (7) and triangular ligament (8) were included to both models as they are needed to maintain tendon alignment and prevent bowstringing during force transmission.

Finally, Fig. 1c shows the tendons from the flexor muscles, *flexor digitorum superficialis* (FDS) *and flexor digitorum profundus*, (FDP). Both the full and trivial models included the same tendons from the flexor muscles. We do not include any connection between the flexor tendons and the extensor mechanism.

Each extensor mechanism model was draped over on the finger bones in an initial configuration according to anatomical data (Garcia-Elias et al., 1991). The model of the bony anatomy included the metacarpal bone, proximal, middle and distal phalanx of the middle finger. The finger joints considered in the model are a metacarpal (MCP; flexion-extension and ad-abduction), proximal interphalangeal (PIP; flexion-extension), and distal interphalangeal (DIP; flexion-extension) joints. The bones were represented as ideal cylinders capped by spheres. The geometric parameters of the cylinders and spheres were based on anatomical surveys (Buchholz et al., 1992; Darowish et al., 2015) to be, respectively: cylinder lengths 64.6 mm, 44.6 mm, 26.3 mm, 17.4 mm; cylinder radii 4.5 mm, 4.0 mm, 3.0 mm, 2.5 mm; the sphere radii 5.0 mm, 5.4 mm, 4 mm for both models.

In addition to bones, five cylinders (a-e in Fig. 1) with smaller radii were included to the model to avoid tendon bowstringing. The cylinder a is perpendicular to the metacarpal bone and replaces a presumed function of the lumbrical muscle pulley (Stack, 1963); the cylinder b is perpendicular to proximal phalanx bone and replaces the presumed function of the protuberances of *p*_1_ head. Cylinders *c, d, e* simulate the annular pulleys that prevent bowstringing of the flexor tendons.

The force of the *extensor digitorum communis* muscle (EDC), ulnar and radial *interosseous* muscle (UI, RI), and lumbrical muscle (LU) were applied to the extensor mechanism model as the input forces. We will note the muscle force values as vector Φ:

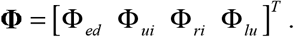

The deformation of the extensor mechanism due to the applied forces and geometric constraints imposed by the bones and the cylinders *a,b* was performed to minimize the overall potential energy (i.e., strain as in (Valero-Cuevas and Lipson, 2004)) of all elastic elements by a gradient algorithm until the equilibrium state was found, as described in (Dogadov et al., 2017).

Once the equilibrium state of the extensor mechanism was found for a set of applied forces, the tendon tensions internal to the extensor mechanism and resulting force at the insertions can be read out. The tensions for each element of the deformed extensor mechanism are found by multiplying its elongation by its stiffness. The forces, transmitted from the extensor mechanism to the bones, including the forces in tendinous insertions and contact forces (the reaction forces created by the tendons overlapping the bones), are used to calculate net joint torques. The torque created by the extensor mechanism were calculated at each kinematic degree of freedom (two for MCP and one each for PIP and PIP). The output fingertip wrench was found as a product of the finger Jacobian inverse transpose, defined by the finger geometry and a posture, with the joint torque vector. This approach is explained in (Valero-Cuevas, 2015; Valero-cuevas et al., 1998).

## RESULTS

Fig. 3 shows the force distribution among the extensor mechanism intercrossing fiber bundles with the posture. The forces in bundles from both side of the finger were similar; therefore, Fig. 3 shows only the forces in the fiber layers from the radial side of the finger. The fiber numbers in Fig. 3 are the same as in Fig. 2. In each posture the extensor mechanism was loaded by four constant muscle forces (UI, EDC, RI, and LU), each of 2.9 N.

**Fig. 3.**
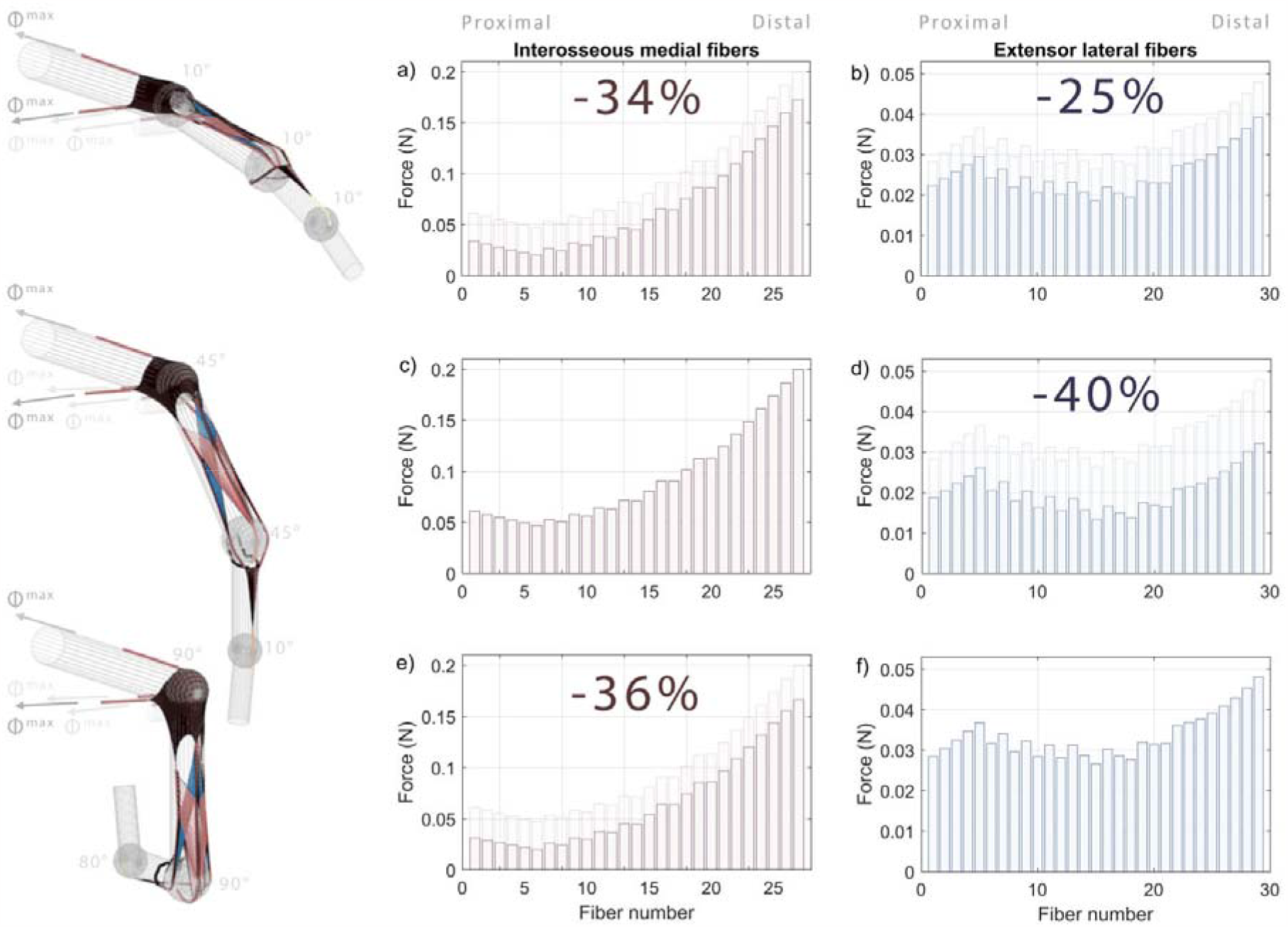
The influence of the posture on the forces forces in intercrossing fiber bundles. Red: interosseous medial fibers; blue: extensor lateral fibers. The first row corresponds to extension (MCP = 10°,PIP = 10°; DIP = 10°), the second row coresponds to mid-flexion (MCP = 45°,PIP = 45°; DIP = 10°), and the third row corresponds to flexion (MCP = 90°,PIP = 90°; DIP = 80°). The input forces were 2.9 N in UI, EDC, RI ans LU muslce for all postures. The maximal value of forces in interosseous medial fibers was attained in mid-flexion and the maximal value in extensor lateral fibers was atteined in full flexion.

It may be seen from the figure that the forces in intercrossing fiber bundles vary with the posture for a constant input force vector. The force in interosseous medial fibers (shown in red) attained the maximal value in mid-flexion posture. The mean force, calculated over all interosseous medial fibers in this posture was 34% higer than in extension and 36% higher than in flexion (0.67 N, 0.66 N and 0.94 N for extension, flexion and mid-flexion сorrespondingly). The forces in extensor lateral fibers arrived to maximal value in flexion posture. The mean force, calculated over all extensor lateral fibers was 25% higher in this posture than in extension and 40% higher than in mid-flexion (0.25 N, 0.20 N and 0.31 N for extension, mid-flexion and flexion correspondingly).

Fig. 4 shows the changes in the feasible tendon force set of the full extensor mechanism model (right column) in comparison with a trivial model (left column). The full-loading state, which was the state when all four extensor muscle forces were equal to Φ^max^, is shown by a circle in each panel.

**Fig. 4.**
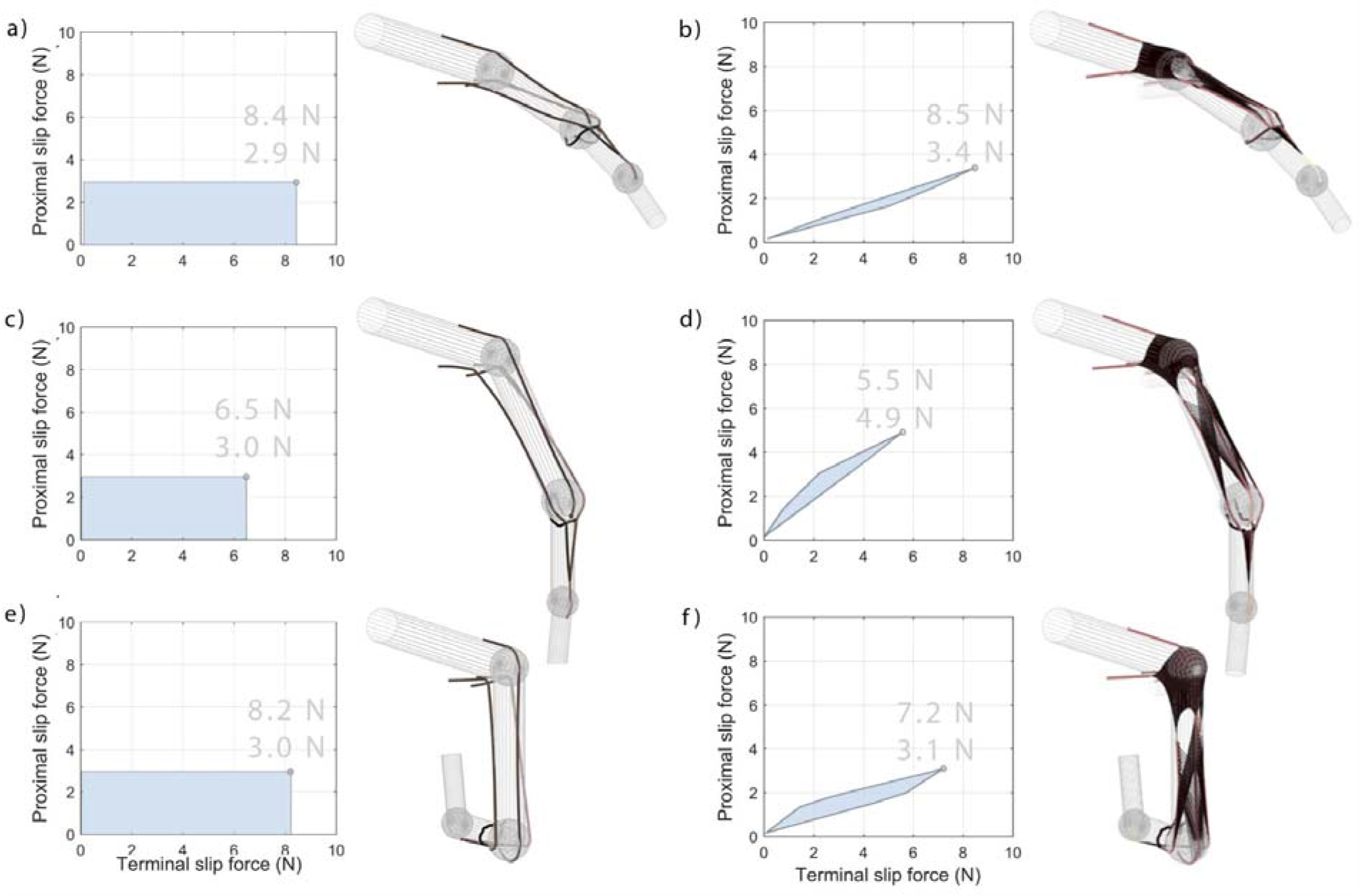
The effect of the posture on feasible tendon force set. Left column corresponds to a trivial extensor mechanism model, right collumn corresponds to a full model. The first row corresponds to extension (MCP = 10°; PIP = 10°; DIP = 10°), the second row coresponds to mid-flexion (MCP = 45°; PIP = 45°; DIP = 10°), and the third row corresponds to flexion posture (MCP = 90°; PIP = 90°; DIP = 80°). The full-loading state, which corresponds to loading of the extensor mechanism models by all four muscles, is shown by a circle in each feasible tendon force set. The proximal and termonal band force values in full-loading sate are comparable for both moodels, but the areas of the feasible tendon force set are smaller for the full model. Also for a full model, the shape and orientation of the feasible tendon force set change with posture

It can be seen from the left column of the image, that the feasible tendon force set of the trivial model had a rectangular shape for all postures. The maximal force in proximal band did not change significantly with posture and was equal to 2.9 N in extension and to 3.0 N in other postures. The maximal force in terminal band was similar in extension and flexion (8.4 N and 8.2 N), but decreased in mid-flexion (6.5 N). This may be explained by the fact that force in terminal tendon is controlled by lateral bands, which are connected by a triangular membrane. Stretching of the triangular membrane in flexion may influence the terminal band force. The ratio between the force in proximal and terminal band of the trivial extensor mechanism model in full-loading state was 0.35 in extension, 0.46 in mid-flexion and 0.37 in flexion.

Contrary to trivial model, the shape, size, and orientation of the feasible tendon force set of the full model strongly change with posture. When the model was loaded by all muscle forces, the force in the proximal band achieved the maximal value in mid-flexion, which was 44% higher than in extension and 58% higher than in flexion (4.9 N, 3.4 N, and 3.1 N for mid-flexion, extension and flexion correspondingly). Contrary to proximal band, the force in terminal band arrived to a minimal value in mid-flexion, which was 35% lower than in extension and 24% lower than in flexion (5.5 N, 8.5 N, and 7.2 N for mid-flexion, extension and flexion correspondingly). As the result of the fact, that the force in proximal and terminal band change differently with posture, the ratio between the force in proximal and terminal band in a full-loading state also varied with posture, and was equal to 0.40 in extension posture, which was the minimal value among all postures. This ratio was maximal in mid-flexion and was equal to 0.89. In flexion, this ratio was equal to 0.43, which is close to the value in extension. It can be also noticed that the area of the feasible tendon forces set for full model was lower than the corresponding areas of the for the trivial model (*e*.*g*. for the extended finger area of the feasible tendon force set for full model was 9% of the feasible tendon force set for trivial model at the same posture).

Fig. 5 shows the effects of the posture on x-y plane projections of the feasible fingertip force set (FFS). The left column corresponds to the trivial extensor mechanism model, the right column to the full extensor mechanism model. The full-loading fingertip force, which was produced by the model when all four extensor muscle forces were equal to Φ^max^, is shown by a circle in each panel. The dark blue area corresponds to the forces created only by the muscles, attached to the extensor mechanism (UI, EDC, RI, and LU) and with no forces in flexor muscles. The light area stands for the forces, created when the flexor muscles were also active (FDS, FDP). For both trivial and full model, shape and orientation of FFS changes with posture were observed.

**Fig. 5.**
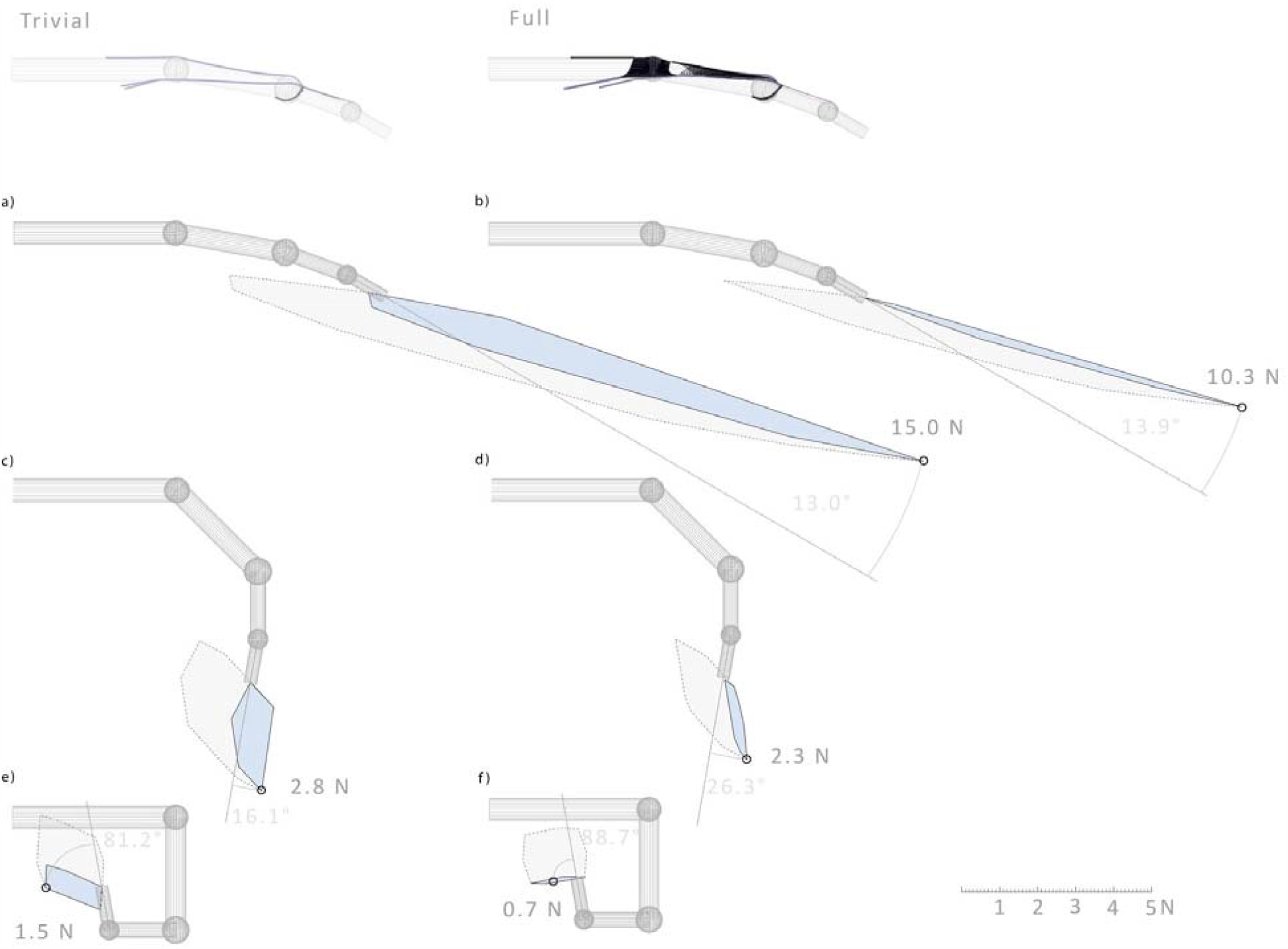
Influence of the posture on x-y plane projection of the feasible force set. Left column corresponds to a trivial extensor mechanism model, right collumn corresponds to a full model. First row corresponds to extension posture (MCP = 10°; PIP ;= 10°; DIP = 10°), second row coresponds to mid-flexion posture (MCP = 45°; PIP = 45°; DIP = 10°), third row corresponds to flexion posture (MCP = 90°; PIP = 90°; DIP = 80°). The area of the feasible force set as well as fingrtip force values in full-loading state are smaller for full extensor mechanism model for all postures. Blue area cooresponds to a subset in a feasible force set produced only by the muscles, attached to the extensor mechanism (UI, EDC, RI, and LU)

The x-y plane projection of the full loading force is lower for the full model than for trivial model for all postures. The full-loading force in full model is 31% lower than in trivial model in extension (10.3 N and 15.0 N correspondingly), 18% lower in mid-flexion (2.3 N and 2.8 N), and 53% lower in flexion (0.7 N and 1.5 N). The angle between the distal phalanx and the xOy-projection of the full-loading force is higher in full model than in trivial one. In extension, the angle in full model and trivial model are 13.9° and 13.0° correspondingly, in mid-flexion the angles are 26.3° and 16.1° and in full flexion the angles are 88.7° and 81.2°. Finally, it can be also noticed from the figure that the area of the FFS of the full extensor mechanism model is lower than the area of the FFS of the trivial model.

## DISCUSSION

We demonstrated that the intercrossing fiber bundles and the extensor hood reduces the area of feasible tendon force set the full extensor mechanism model, which contain the intercrossing fibers and the extensor hood, is lower than the areas of feasible tendon force set and FFSs, produced by the trivial model, in which there are no connections between the medial and lateral bands. This area increases due to the fact that the trivial extensor mechanism model enables the independent control of the forces in the proximal and terminal band. However, in the case of the full model of the extensor mechanism, these forces are naturally coupled.

Secondly, we have shown that the bundles of intercrossing fiber can modify the force distribution according to posture. This may indicate that the nervous system has to modulate the sharing in involved muscle and intensity according to the finger posture in order to produce the wanted fingertip force. This may imply that there exists a link between the passive adaptations of the extensor mechanisms and the active modulation of the muscle recruitments for useful fingertip tasks, such as grasping objects (Wei et al., 2022), writing (Gerth and Festman, 2023), or playing musical instrument (Furuya et al., 2011).

The analyzed full model has several limitations. Firstly, the model topology oversimplifies the real extensor mechanism anatomy. Over MCP joint the extensor mechanism was represented only by the extensor hood. However, the metacarpophalangeal fibrous griddle, or sagittal band, which connect the extensor tendons to the deep transverse intermetacarpal ligament and capsular join (Zancolli, 1979) was not taken into account. Moreover, no attachments of the extensor mechanism at the base of the proximal phalanx were taken into account. Secondly, the bones were modeled as cylinders with spheres corresponding to the joints.

This study is limited in that it does not include all other muscles acting on the finger, but this work enables future work to understand the function of the human fingers that considers their complex anatomy in more detail.

In addition, this work only considered force transmission by the trivial model, but does not consider other important biomechanical consequences of it. First and foremost is the need to maintain and regulate the tendon path as the finger changes posture, where the “unsupported” trivial tendons may slide, bowstring, cause rapid changes in moment arms and even cause tendinitis or scaring during their unguided sliding movement. In our model, the path of the tendons in the trivial model was enforced arbitrarily. From this perspective, the extensor mechanism may server to retain force transmission while also serving as a support and guiding structure, much like the annular bands and sesamoids in other tendons.

And secondly, there are other considerations in addition to tendon force and joint roque production. Recent work has suggested that tendon force transmission is important for other important aspects of function such as stability during force production (Sharma and Venkadesan, 2022). Similarly, producing slow finger movements very likely depend more on managing the internal strain energy of the system and not second-order rigid-body dynamics driven by joint torques or muscle forces (Babikian et al., 2016).

As such, the evolutionary pressures for the formation of the extensor mechanism may not be strictly limited to force transmission. That is, the extensor mechanisms may have been a multi-factorial evolutionary adaptation that also allows for stability and accurate slow movements with the fingertips that gave human-primates a competitive advantage for effective manipulation capabilities.

## ACKNOWLEDGEMENTS

The work was supported by IDEX scholarship for international mobility. The author acknowledge Vishweshwer Shastri (USC) and Gelu Ionescu (GIPSA-Lab) for their assistance with the programming of the extensor mechanism simulator.

## Notes

### Competing Interest Statement

The authors have declared no competing interest.

